# The bacterial microbiome of symbiotic and menthol-bleached polyps of long-term aquarium-reared *Galaxea fascicularis*

**DOI:** 10.1101/2023.08.23.554380

**Authors:** Giulia Puntin, Jane C. Y. Wong, Till Röthig, David M. Baker, Michael Sweet, Maren Ziegler

## Abstract

Coral reefs support the livelihood of half a billion people but are at high risk of collapse due to the vulnerability of corals to climate change and local anthropogenic stressors. While understanding coral functioning is essential to guide conservation efforts, research is challenged by the complex nature of corals. They exist as metaorganisms (holobionts), constituted by the association between the (coral) animal host, its obligate endosymbiotic algae (Symbiodiniaceae), and other microorganisms comprising bacteria, viruses, archaea, fungi and other protists. Researchers therefore increasingly turn to model organisms to unravel holobiont complexity, dynamics, and how these determine the health and fitness of corals. The coral Galaxea fascicularis is an emerging model organism for coral symbiosis research with demonstrated suitability to aquarium rearing and reproduction, and to manipulation of the host-Symbiodiniaceae symbiosis. However, little is known about the response of the *G. fascicularis* microbiome to menthol bleaching—the experimental removal of the Symbiodiniaceae which represents the first step in coral-algal symbiosis manipulation. For this, we characterized the bacterial microbiome of symbiotic and menthol-bleached *G. fascicularis* originating from the Red Sea and South China Sea (Hong Kong) that were long-term aquarium-reared in separate facilities. We found that the coral-associated microbiomes were composed of relatively few bacterial taxa (10-78 ASVs). Symbiotic polyps (clonal replicates) from the same colony had similar microbiomes, which were distinct from those of other colonies despite co-culturing in shared aquaria. A pattern of seemingly differential response of the bacterial microbiome to menthol bleaching between the two facilities emerged, warranting further investigation into the role of rearing conditions. Nevertheless, the changes in community composition overall appeared to be stochastic suggesting a dysbiotic state. Considering the importance of bleaching treatment of captive corals for symbiosis research, our results—although preliminary—contribute fundamental knowledge for the development of the Galaxea model for coral symbiosis research.

## Introduction

By the end of this century, the livelihood of more than half a billion people will largely depend on the ability of corals to cope with changing ocean conditions (Hughes et al. 2017). Corals are vulnerable to climate change, pollution, and overfishing, which have caused the loss of half of the world’s coral cover since the 1950s (Eddy et al. 2021; Eakin et al. 2022). As such, the reef structures they build and the multi-billion dollar ecosystem services they provide are at high risk of collapse (Costanza et al. 2014; van Hooidonk et al. 2016). Considering the high stakes, understanding coral functioning is essential to predict future scenarios and guide management and conservation efforts.

Corals exist as metaorganisms, or so-called holobionts (Rohwer et al. 2002), where complexity hinders our ability to unravel coral functioning and how it ultimately affects coral physiology and ecology (Rosenberg et al. 2007; Jaspers et al. 2019). Indeed, the coral holobiont comprises the animal host, obligate intracellular algal symbionts (Symbiodiniaceae), a rich and diverse bacterial community, together with other microorganisms such as archaea, fungi, viruses, and protists (Bourne et al. 2016; Pogoreutz et al. 2020). Even richer than their taxonomy is the potential diversity of relationships and interactions among members, and how these could contribute to holobiont health and resilience (Thompson et al. 2015; Pogoreutz et al. 2020).

Besides the coral host, the best understood member of the coral holobiont is the algal symbiont, as its photosynthesis-derived energy fuels the calcification process that builds reefs and allows corals to thrive in nutrient-poor waters (Muscatine and Porter 1977; Muscatine 1990). Bacteria are also involved in nutrient cycling and metabolism (Robbins et al. 2019; Tandon et al. 2020), as well as other essential physiological processes such as development (Webster et al. 2004; Tebben et al. 2015) and immunity (Certner and Vollmer 2018; Miura et al. 2019). Interestingly, bacteria also mediate host-Symbiodiniaceae dynamics through the mitigation of thermal and light stress on the algal symbiont (Motone et al. 2020; Connelly et al. 2022), and they produce antimicrobial agents from an organosulfur compound released by Symbiodiniaceae (Raina et al. 2016, 2017). A growing body of knowledge suggests a central role of complex multipartite interactions between bacteria, Symbiodiniaceae, and the coral host in nutrition, health and fitness (reviewed in Matthews et al. 2020). However, given the intricacy of players’ diversity and their metabolic capacities, linking partner identity to function and holobiont phenotype proves particularly challenging.

Model organisms can help unravel holobiont complexity through manipulation. Comparison of different host-Symbiodiniaceae combinations in the sea anemone Aiptasia revealed that heat-tolerance of the symbiont is not linearly transferred to the host (Chakravarti et al. 2017; Gabay et al. 2019; Herrera et al. 2020). Thus complex mechanisms where holobiont properties cannot be predicted as the “sum of its parts” require a more holistic approach (Goulet et al. 2020). Such empirical testing of multi-partner interactions relies on the ability to study the holobiont upon experimental manipulation, namely by removing and/or adding members (Jaspers et al. 2019). Most of such studies in the field were conducted with non-calcifying species such as Aiptasia or the hydroid polyp *Hydra*, which are well established, tractable model organisms (Weis et al. 2008; Galliot 2012). Yet, while these models have proven instrumental for breakthrough discoveries in the cnidarian symbiosis field (e.g., Weis 2008; Murillo-Rincon et al. 2017; Pietschke et al. 2017; Gabay et al. 2018), they lack key features of reef-building corals such as calcification and the obligate endosymbiosis necessary to understand the ecology of coral functioning. To include these critical features, research with corals is irreplaceable (Puntin et al. 2022b).

The reef-building coral *Galaxea fascicularis* (Linnaeus, 1767) has been proposed as a model species for coral symbiosis research (Puntin et al. 2022a). This species is well represented in the literature, featured in studies that characterize the coral gastric cavity (Agostini et al. 2012; Zhou et al. 2020) and the calcification processes (Al-Horani et al. 2003, 2005, 2007), among others (Ferrier-Pagès et al. 1998; Niu et al. 2016; Miura et al. 2019). Introducing *G. fascicularis* as a model system, we previously demonstrated its ease of rearing in simplified systems (closed, small volume), compatibility with *ex-situ* reproduction, effective removal of the algal symbiont through menthol bleaching, and subsequent reestablishment of the symbiosis with both cultured and environmental Symbiodiniaceae in adult individuals (Puntin et al. 2022a). This demonstrated the potential to experimentally produce a variety of coral-Symbiodiniaceae combinations to study symbiosis functioning and partner compatibility in a true reef-building coral. While recent coral probiotic approaches rapidly expand our knowledge on the functions of the bacterial fraction (Rosado et al. 2019; Peixoto et al. 2021), untangling the complexity in the holobiont requires detailed knowledge of the interrelationships that consider all partners.

One of the main knowledge gaps to advance mechanistic symbiosis research in the Galaxea model system is the effect of menthol bleaching on the remaining coral microbiome. Menthol is becoming increasingly common in manipulative experiments due to its efficacy in removing Symbiodiniaceae while causing virtually no host mortality (Wang et al. 2012; Matthews et al. 2015; Puntin et al. 2022a). To date, menthol bleaching has been used with a range of symbiotic cnidarians, including jellyfish (Röthig et al. 2021), anemones (Matthews et al. 2015; Dani et al. 2016), corallimorpharia (Lin et al. 2019), and nine species of reef-building corals (Wang et al. 2012, 2019; Puntin et al. 2022a; Scharfenstein et al. 2022; Chan et al. 2023). Yet, its impact on their bacterial fraction remains unknown. Menthol is known to have antimicrobial activity against several human pathogens (Trombetta et al. 2005; Mahzoon et al. 2022) and can select against certain bacteria, thus impacting microbial community composition, as seen in other contexts (Chopyk et al. 2017). To address this knowledge gap, we characterized the bacterial microbiome of symbiotic and menthol-bleached *G. fascicularis* polyps from the central Red Sea and the South China Sea that were maintained in two separate facilities for several months.

## Materials and Methods

### Coral collection and long-term aquarium rearing

Colonies of *Galaxea fascicularis* were collected from two locations: the Red Sea (hereafter referred as “Red Sea”) and Hong Kong in the South China Sea (hereafter referred as “Hong Kong”). Red Sea colonies (n = 3) were collected from the central Saudi Arabian Red Sea at “Al Fahal” reef (N 22 ° 18.324’ E 38 ° 57.930’), at 9-13 m depth in March 2019 (CITES permit 19-SA-000096-PD) and transported to the Ocean2100 aquarium facility at Justus Liebig University Giessen (Germany) where they were maintained in a 7,000 L closed aquarium system composed of several tanks (100-265 L) including a technical tank fitted with protein skimmer, active charcoal filter, phosphate adsorber, algal refugium (*Chaetomorpha* sp.) with reverse light cycle, and calcium reactor (Schubert and Wilke 2018). In the aquarium system, light was provided by white and blue fluorescent lamps with a light:dark cycle of 12:12 h at 130-160 µmol photons m^−2^ s^−1^ to approximate light conditions at the collection site (Ziegler et al. 2015). Salinity was maintained around 35 and temperature at 26 °C. Colonies were fed daily with a combination of frozen copepods, Artemia, krill, and Mysis. Hong Kong colonies (n = 2) were collected from ≤ 5 m depth from Crescent Island (N 22° 31’ 51.035”, E 114° 18’ 53.888) in June 2019 and transported to the University of Hong Kong (HKU), where they were maintained in a 500-L aquarium equipped with a filtration system and protein skimmer, and fed daily with Reef-Roids (Polyplab) and frozen artemia. Light intensity, salinity, and temperature conditions were consistent with those maintained in the Ocean2100 facility.

### Menthol bleaching

At both locations, individual (clonal) polyps were mechanically isolated from their colony, mounted on coral glue (Red Sea colonies, JLU-Ocean2100; Grotech, Cora-Fix SuperFast) or attached to small ceramic tiles (Hong Kong colonies, HKU; Aron Alpha, GEL-10) (Fig. 1). After 10-14 days of healing, polyps were randomly divided between a ‘symbiotic’ and a ‘bleached’ group at each location. Both groups were maintained under the same conditions until healed, then the ‘bleached’ group was treated with menthol to chemically induce bleaching. Menthol treatment started after approximately seven (Red Sea) and three (Hong Kong) months of captivity, and it was replicated at the two facilities following a protocol modified from Wang et al. (2012). Specifically, three days treatment in 0.38 mM menthol solution in filtered (1.2 µm) artificial seawater (FASW) was followed by one day of rest and another day of menthol treatment. Menthol incubations lasted 8 h during the light period.

**Figure 1.**
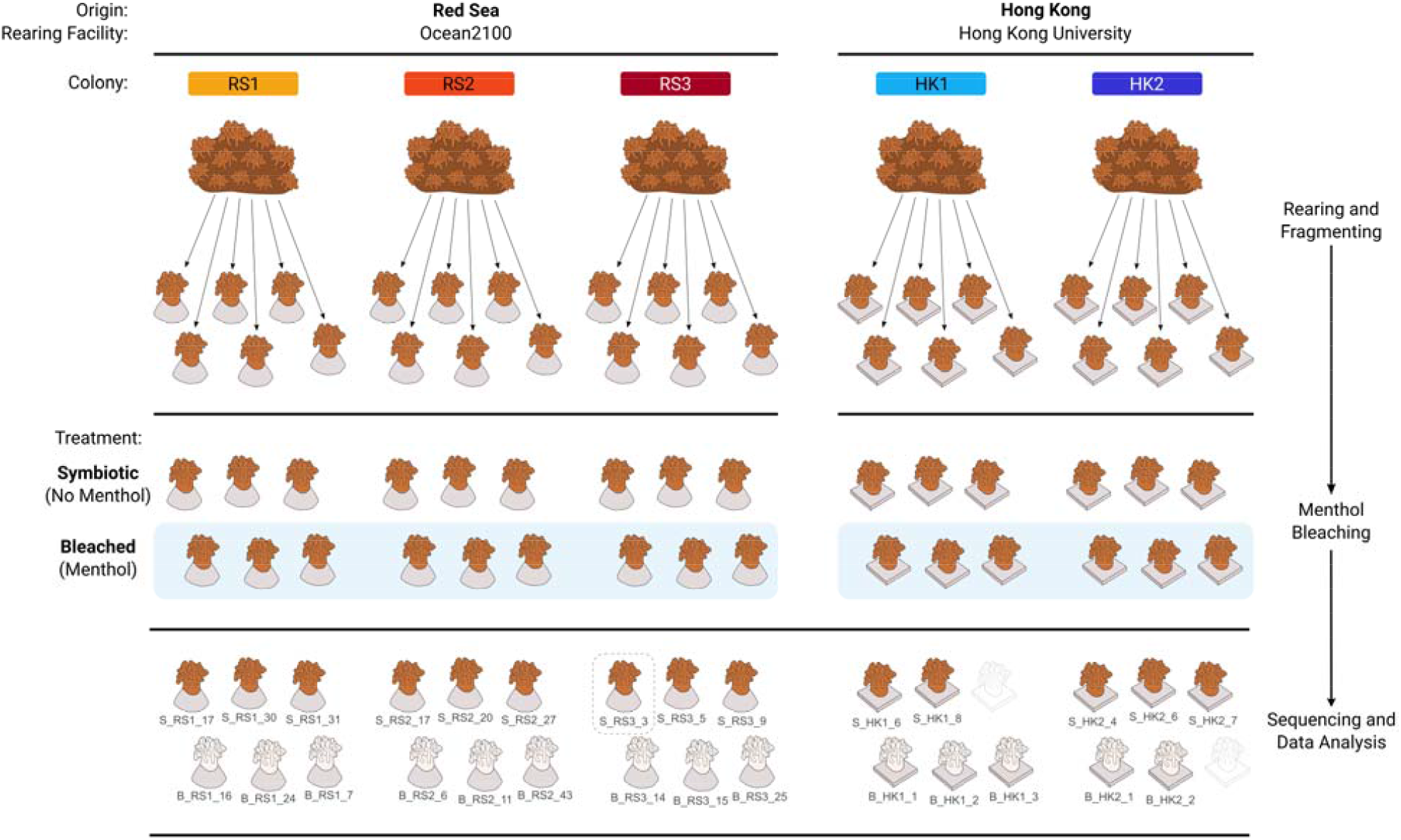
Visual summary of the experimental design and data processing. Two samples (outlined polyps) were excluded from processing due to damage during shipping, while one sample (dashed box) was omitted after rarefaction for alpha and beta diversity analysis owing to low sequencing depth.

Bleaching was assessed in the Red Sea polyps by visual inspection under a fluorescence stereomicroscope (Leica MZ16 F) 10 days after the menthol treatment, when algal cells were not detectable in any polyp. At the same time point, Hong Kong polyps also appeared fully bleached under microscopic inspection (Olympus Optical, mod. CHK at 400×).

### Post-bleaching rearing conditions

To prevent coral from Symbiodiniaceae exposure and symbiosis re-establishment, all polyps were kept in simplified (see below for details) systems with FASW (1.2 µm) after the menthol bleaching treatment. Here, polyps were fed daily with one small frozen adult Artemia each, followed by partial (∼10 %) water change after 2-3 h. At both facilities, temperature, light, and salinity were maintained consistent with the long-term rearing conditions, while the setups differed. Specifically, at the Ocean2100 facility, the polyps were distributed among eight 5-L glass tanks (20 cm × 30 cm) (four per treatment), each equipped with a small pump (Resun SP-500) in a temperature-controlled water bath. At HKU, symbiotic and bleached polyps were maintained in separate 600 ml glass jars, each holding ∼6 polyps and equipped with magnetic stir bars for water flow inside a Plant Growth Chamber (Panasonic MLR-352H-PA).

### Sampling for microbial analysis

16S rRNA amplicon sequencing was employed to characterize the bacterial communities of symbiotic and menthol-bleached polyps of *G. fascicularis* colonies from the two geographic locations (Fig. 1).

At both locations, three polyps per colony (5 colonies: RS1, RS2, RS3, HK1, HK2) per state (2 states: symbiotic, bleached) were sampled on the 13^th^ day after the menthol treatment when they visually appeared completely bleached and otherwise healthy (n = 15 bleached and 15 symbiotic polyps). The polyps were rinsed with seawater, separated from the substrate, placed in sterile tubes, and stored at −80 °C. For transport, polyps were stored in RNAlater (R0901-100ML, Sigma-Aldrich, Hong Kong S.A.R.). Two samples from Hong Kong (1 symbiotic and 1 bleached) were damaged during shipping and therefore excluded from processing. All samples were processed together for DNA extraction (University of Derby’s Aquatic Research Facility, UK) and subsequently sequenced in the same sequencing run (Bart’s and the London Genome Centre, Queen Mary, University of London).

### Bacterial community analysis

DNA was extracted using the Qiagen DNeasy 96 Blood & Tissue kit from crushed polyps (containing skeletal and tissue material) with about 10 mg of coral tissue per sample as starting material. Extractions followed the user manual with centrifugations at working steps 10, 12, and 16 performed at 1500 g at doubled centrifugation times. Extracted DNA was sent to the sequencing facility for quality control, PCR, library preparation, and pair-end sequencing with Illumina MiSeq platform v3 (2 × 300 bp). The 16S rRNA gene region V5/V6 was amplified using the primers 784F and 1061R (Andersson et al. 2008). A contamination control consisting of pure RNAlater buffer was included in all steps.

Bacterial sequencing data were processed in Qiime2 (v.2021.11, Bolyen et al. 2019) and analyzed in R (v.4.1.0, R Core Team 2021). After primer removal, forward and reverse reads were truncated to 232 and 234 nt respectively, paired, dereplicated, quality checked, cleaned, and clustered to amplicon sequence variants (ASVs) using the denoise-paired method in DADA2 (Callahan et al. 2016). This resulted in a total of 138,620 sequences and 547 ASVs. ASVs were taxonomically assigned using a weighted classifier trained against the SILVA 138 database (99 % clustering, full length) (Yilmaz et al. 2013) with the classify-sklearn method from ‘q2-feature-classifier’ plug-in (Bokulich et al. 2018; Kaehler et al. 2019; Robeson et al. 2020). Then, sequences assigned to “mitochondria”, “chloroplast”, “Archaea”, “Eukaryota”, or “unknown” at the phylum level were removed. Sequences found in the control sample were considered potential lab contaminants and evaluated based on presence/absence across the coral samples and habitat description (e.g., known contaminants) of the closest BLASTn matches (GenBank) (for full details see https://zenodo.org/record/10551928, “01_find_contaminants.R”). This led to the removal of five ASVs and their 18,990 sequences, and resulted in a final data set of 112,789 sequences and 515 ASVs across 28 samples (after excluding the contamination control).

Rarefaction and alpha and beta diversity calculations were performed with the R package ‘phyloseq’ (v.1.38.0, McMurdie and Holmes 2013), ‘metagMisc’ (v.0.0.4, Mikryukov 2023) and ‘btools’ (v.0.0.1, Battaglia 2022). Samples were rarefied to 2,690 sequences (based on 1,000 iterations of random subsampling without replacement), which caused the exclusion of one sample (symbiotic colony “RS3”) due to low sequencing depth. Rarefaction curves showed that for most samples the number of ASVs plateaued before the rarefaction depth indicating that most of the diversity was captured and retained after rarefaction (Fig. S1). Alpha diversity was estimated through multiple indices chosen for their complementarity and comparability with previous studies (observed richness, Chao1, Shannon diversity, Simpson diversity, Pielou’s evenness, and Faith’s phylogenetic diversity). Differences in alpha diversity between symbiotic and bleached individuals were tested with t-tests or Mann-Whitney U tests depending on data distribution and variance. Beta diversity based on Bray-Curtis distances was visualized with non-metric multidimensional scaling (nMDS). The differential relative abundance of bacterial taxa between symbiotic states (symbiotic, bleached) was tested at the ASV and family level across the whole data set and by geographic origin (Red Sea, Hong Kong) using the Mann-Whitney U test with Benjamini-Hochman correction for multiple comparison. Relative abundance bubble plots of bacterial community composition at bacterial family level and of core ASVs (occur in all groups by state and geographic origin), and the UpSet plot (Lex et al. 2014) were generated from non-rarefied data. The UpSet plot was created with the package ‘UpSetR’ (v.1.4.0, Conway et al. 2017). All other plots were created in ‘ggplot2’ (v.3.3.5, Wickham 2016).

## Results

### Microbial diversity and richness were unaffected by menthol bleaching

When considering all colonies together, alpha diversity remained similar between symbiotic and menthol-bleached samples across all diversity and richness indices tested (incl. observed richness, Chao1, Shannon diversity, Simpson diversity, Pielou’s evenness, and Faith’s phylogenetic diversity, see Table S1, S2), and regardless of their origin (P_t-test_ or P_Mann-Whitney U-test_ > 0.05; Fig. 2A). When considering each individual colony, alpha diversity was remarkably and consistently low in symbiotic RS1 polyps, while in their bleached counterparts it was in range with the other colonies (Fig. 2A, Fig. S2, Tab. S2). Within-colony difference between symbiotic and bleached polyps could only be tested for RS1 and RS2, and it was significant in RS1 across all alpha diversity indices tested (P_Welch_ < 0.05, Tab. S3).

**Figure 2.**
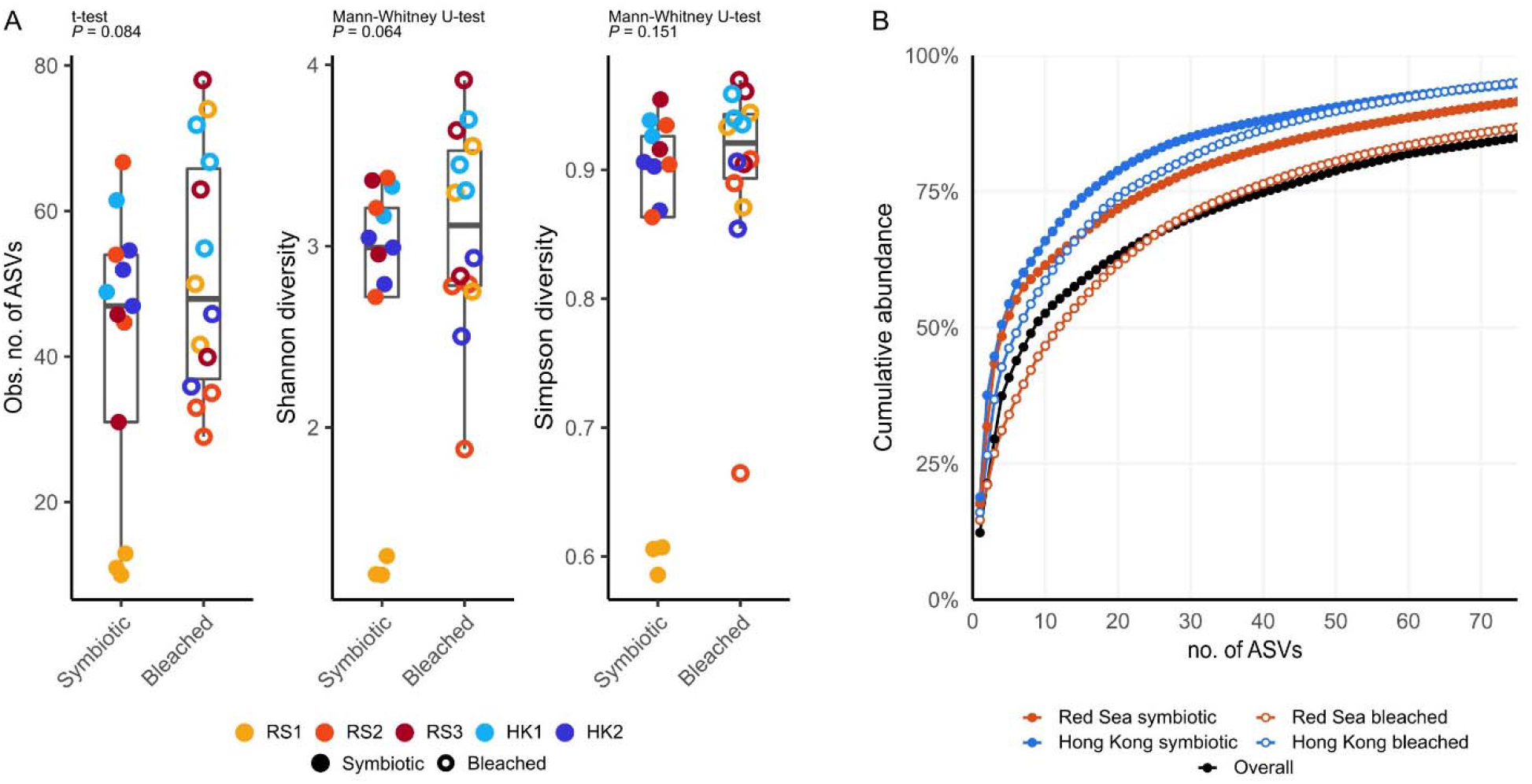
Diversity and evenness of the bacterial communities of symbiotic and menthol-bleached polyps of *Galaxea fascicularis*. (A) Comparison of observed number of ASVs, and Shannon and Simpson diversity index between symbiotic and bleached samples; (B) ASV accumulation curve for the whole data set, and separated by symbiotic state and geographic origin.

### Bacterial communities were generally uneven

A small number of ASVs dominated the bacterial communities, where the 3 and 9 most abundant ASVs accounted for > 25 % and > 50 % of the total number of sequences, respectively (Fig. 2B). Evenness was on average lower among the symbiotic polyps. Specifically, in the symbiotic samples, the 5 (for Red Sea) and 4 (for Hong Kong) most abundant ASVs account for > 50 % of total reads, while in the bleached samples it took 7 (for Red Sea) and 12 (for Hong Kong) ASVs to pass the 50 % relative abundance threshold. Of note, microbiomes were made of a small number of bacterial taxa, ranging from 10 to 78 ASVs, and with symbiotic RS1 polyps having the smallest number of ASVs (10, 11, and 13 respectively).

### Menthol-bleaching seemed to elicit stochastic changes in the microbial communities

Changes in community composition between symbiotic states showed different trends across the two facilities, however dissimilarity was generally higher in menthol bleached polyps. While symbiotic polyps clustered by colony for both Red Sea and Hong Kong (indicating similar microbial communities, Fig.3A), bleached polyps showed no such clear grouping (Red Sea, Fig.3C) or larger scattering compared to symbiotic polyps (Hong Kong, Fig.3D), and collectively had a significantly higher within-colony dissimilarity (P_Mann-Whitney_ = 0.0001, Fig. 3A,B, Fig. S3), seemingly indicating random changes in the communities of the menthol-bleached polyps.

**Figure 3.**
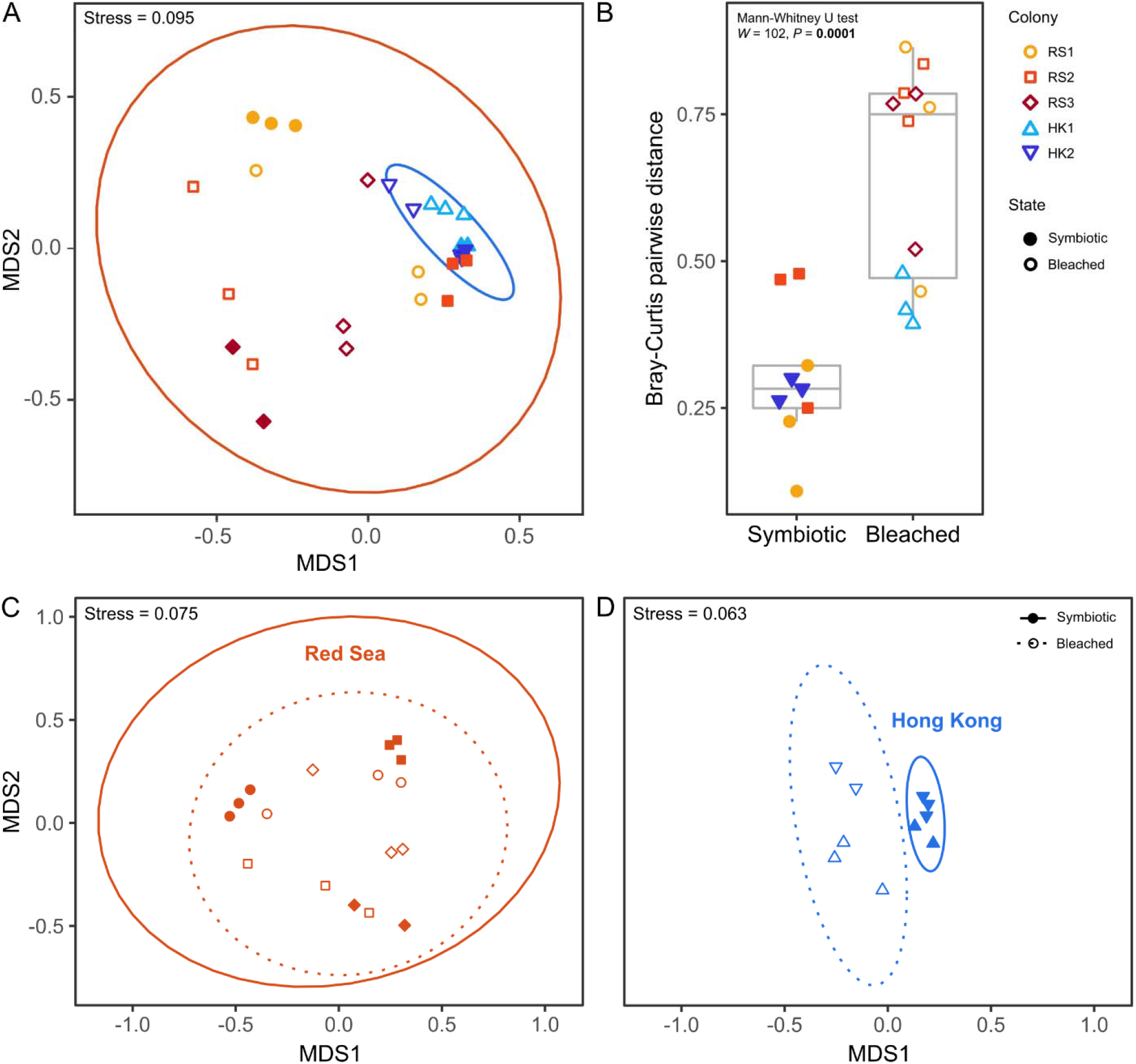
Microbial community structure of *Galaxea fascicularis* from the Red Sea and Hong Kong. Non-metric multi-dimensional scaling (nMDS) plot of bacterial community composition based on Bray-Curtis dissimilarity for all polyps (A), Red Sea polyps (C), and Hong Kong polyps (D), and comparison of pairwise dissimilarity between symbiotic and bleached polyps of the same colony (only groups with *n* = 3 considered, but comparable results were found considering groups with *n* < 3, see Fig. S3) (B). Ellipses = 95 % confidence intervals (A, C, D); colors (A, D) or shapes (C, D) denote colony identity, filled symbols = symbiotic polyps, hollow symbols = bleached polyps.

**Figure 4.**
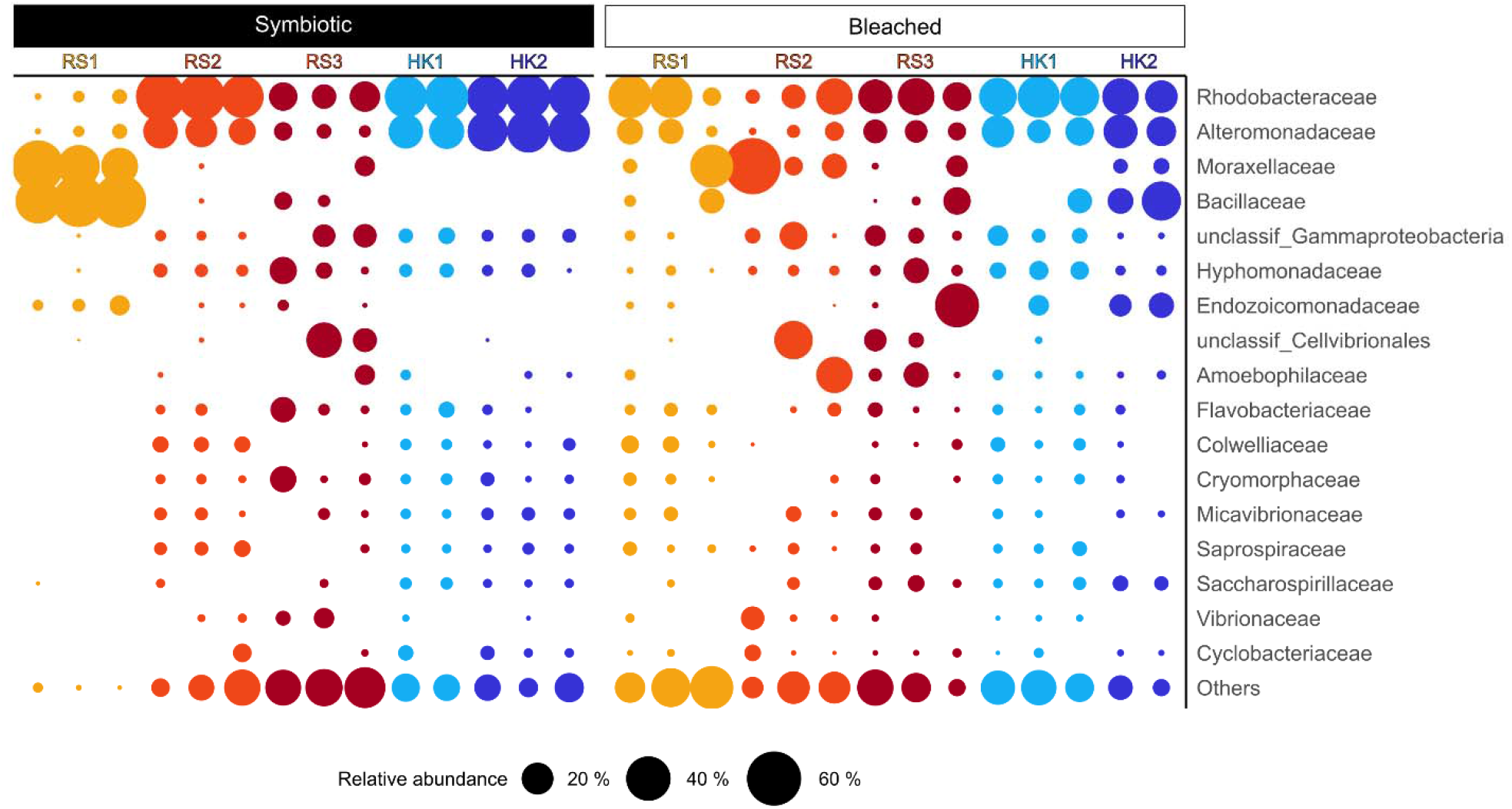
Relative abundance of bacterial families in symbiotic and bleached *Galaxea fascicularis* polyps from the Red Sea and Hong Kong. Bubble size is proportional to the relative abundance per polyp of the 17 most abundant families, which account for 83.7 % of the total number of reads; all other 121 families that had a relative abundance < 1 % are grouped as ‘Others’.

Additionally, no bacterial taxa (neither at ASV nor at bacterial familiy level) showed significantly different relative abundance between symbiotic and bleached samples, neither across the whole data set nor by geographic origin (i.e., considering Red Sea and Hong Kong samples separately) (Mann-Whitney U test with Benjamini-Hochman correction, all P > 0.05). This includes bacteria previously reported to be associated with Symbiodiniaceae cultures (Fig. S4; Supplementary Materials and Methods).

### Three bacterial families dominated most microbial communities

A total of 138 bacterial families were found across the 28 sampled polyps. Of these, 104 bacterial families occurred in symbiotic (74 in Red Sea, 66 in Hong Kong) and 109 in bleached (89 in Red Sea, 69 in Hong Kong) polyps. The three most abundant families *Rhodobacteraceae* (25.8 %), *Alteromonadaceae* (14.6 %), and *Moraxellaceae* (10.1 %) together represented > 50 % of all sequences, and the 11 most abundant families represented > 75 % of all sequences. Symbiotic colonies were largely dominated by members of the two bacterial families *Rhodobacteraceae* and *Alteromonadaceae*. Except for the two Red Sea colonies RS1 and RS3 that were dominated by *Moraxellaceae* and *Bacillaceae*, or that showed no consistently dominant family across individual polyps, respectively.

Bleached polyps were dominated by varying bacterial families, with inconsistent patterns between and within colonies and regions. In addition to the dominant families in symbiotic colonies, *Endozoicomonadaceae*, unclassified *Cellvibrionales*, and *Amoebophilaceae* became dominant in some bleached polyps. While members of the family *Endozoicomonadaceae* were the most abundant fraction in one bleached polyp (39 %; RS3), they were only present in 15 of the 28 polyps (7 symbiotic, 8 bleached), and with low relative abundance (mean 2.2 % in symbiotic, and 8.9 % in bleached).

### 28 core ASVs were shared by all coral colonies and symbiotic states

Altogether, there were more exclusive than shared ASVs between experimental groups (Fig. 5A). Specifically, 33.4 % and 52.9 % of ASVs were exclusively found in symbiotic or bleached samples, respectively, while 5.4 % of ASVs (28) were found in all groups (symbiotic state and geographic origin) and were considered core taxa. Although no ASV was found in all samples (nor in all samples of the same experimental group, Tab. S4), three ASVs occurred in ≥ 70 % of samples, and five additional ASVs occurred in ≥ 60 % of samples (Fig. 5B). The sequences of these eight ASVs, which also corresponded to the most abundant of the 28 core ASVs, were BLAST-searched against the NCBI Nucleotide collection (Tab. 1), which revealed that most of the matching sequences were from samples associated with marine environments and/or organisms. Among these, the most abundant bacterial sequences belonged to the genera *Alteromonas, Ruegeria*, and *Nautella* (Fig. 5B, Tab. 1). Interestingly, two ASVs (ASV_001 and ASV_006), both assigned to the genus Ruegeria, matched with strains isolated from aquarium-reared *Galaxea fascicularis* from the South China Sea (collected in Hainan Island, China, Zhou et al. 2020) and Japan (Miura et al. 2019). Further, three ASVs (ASV_006, ASV_018, ASV_020) matched with bacteria identified in Acropora spp. and *Pocillopora* spp. from the central Red Sea, amplified with the same primer set used for this study (Tab. 1).

**Table 1.** Summary of NCBI BLASTn matches for the eight most common core ASVs (occur in both symbiotic and bleached samples from both Red Sea and Hong Kong, and in > 60 % of all samples) in order of abundance. For brevity, only the top three matches are reported (100 % query cover) with information on isolation source and location, where coral species are highlighted in bold. Taxonomy is reported as the lowest taxonomic level assigned by the Qiime2 classifier.

| Taxonomy | ASV no. | GenBank<br>Accession no. | Identity<br>(%) | Isolation source (location) |
| --- | --- | --- | --- | --- |
| *Ruegeria* | ASV_001 | MK967091 | 100.00 | Jellyfish, *Aurelia aurita* polyp (Kiel Bight, Baltic Sea) |
|  |  | MT187950 | 100.00 | ***Galaxea fascicularis*** (Hainan, South China Sea) |
|  |  | MT187656 | 100.00 | ***Galaxea fascicularis*** (Hainan, South China Sea) |
| *Alteromonas* | ASV_003 | MT525287 | 100.00 | Marine sediment (South Korea) |
|  |  | MK967157 | 100.00 | Artificial Seawater (Kiel Bight, Baltic Sea) |
|  |  | MT515801 | 100.00 | Marine sediment (South Korea) |
| *Nautella* | ASV_005 | LC543501 | 100.00 | Coastal seawater (Japan) |
|  |  | MH556771 | 100.00 | Ciliate, *Euplotes vannus* in Yantai Institute of Coastal Zone Research (Shandong, China) |
|  |  | MK801649 | 100.00 | Unspecified coral from South China Sea (Guangdong, China) |
| *Ruegeria* | ASV_006 | MK736137 | 100.00 | ***Acropora hemprichii*** (Central Red Sea) |
|  |  | MK736054 | 100.00 | ***Pocillopora verrucosa*** (Central Red Sea) |
|  |  | MH807604 | 100.00 | ***Galaxea fascicularis*** (Aquarium, Japan) |
| *Alteromonas* | ASV_008 | MT472680 | 100.00 | Shrimp gut, *Penaeus vannamei* (India) |
|  |  | MN704564 | 100.00 | Undescribed source at marine molecular ecology lab (Zhejiang, China) |
|  |  | MH556744 | 100.00 | Ciliate, *Euplotes vannus* in Yantai Institute of Coastal Zone Research (Shandong, China) |
| unclassified<br>*Gammaproteobacteria* | ASV_017 | AB294979 | 99.61 | Microbial mat at shallow submarine hot spring (Okinawa, Japan) |
|  |  | KU629853 | 99.23 | Seaweed, *Asparagopsis* sp. (Portugal) |
|  |  | GU061986 | 99.23 | Oceanic water (South China Sea) |
| unclassified<br>*Cryomorphaceae* | ASV_018 | KY374050 | 100.00 | ***Acropora hyacinthus*** (Central Red Sea) |
|  |  | KF185498 | 99.60 | Marine snow (Adriatic Sea) |
|  |  | KU648372 | 95.65 | Seawater (Gullmarsfjorden, Sweden) |
| unclassified<br>*Hyphomonadaceae* | ASV_020 | KY455482 | 100.00 | ***Acropora hemprichii*** (Central Red Sea) |
|  |  | KY373570 | 100.00 | ***Acropora hyacinthus*** (Central Red Sea) |
|  |  | CP017718 | 100.00 | Algal culture, coral reef substrate (Palmyra Atoll, Pacific Ocean) |

**Figure 5.**
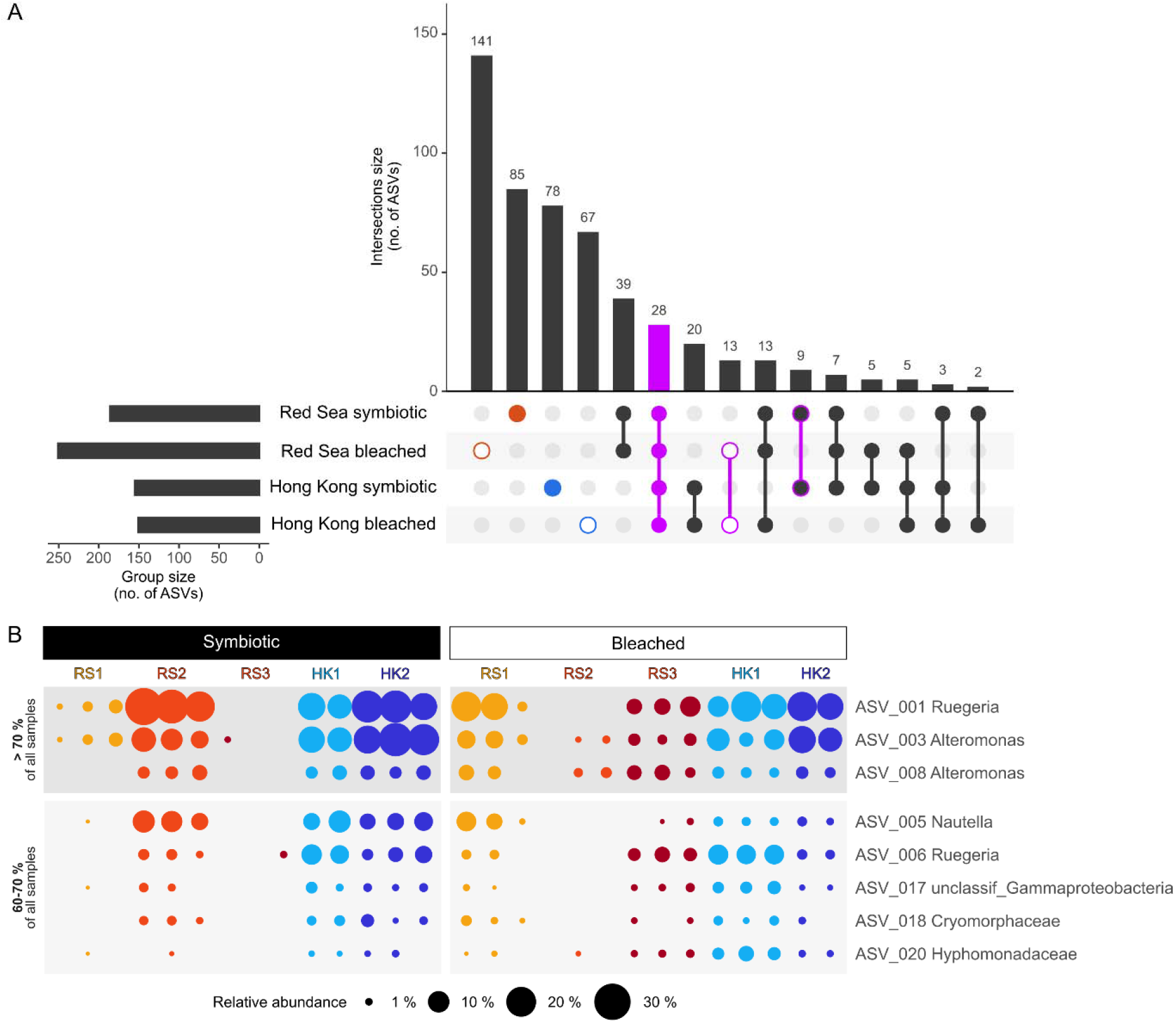
Overview of total and core ASVs in *Galaxea fascicularis* symbiotic and menthol-bleached polyps from the Red Sea and Hong Kong. (A) UpSet diagram showing the number of observed ASVs per group and per intersection. (B) Relative abundance of core bacterial ASVs by host origin and symbiotic state, considering only ASVs present in ≥ 70 % (top-darker grey block) and between 60 % and 70 % of samples (bottom-lighter grey block); bubble size is proportional to the relative abundance of ASVs in each sample.

## Discussion

We employed 16S rRNA gene amplicon sequencing to characterize the bacterial communities of symbiotic and menthol-bleached G. *fascicularis* polyps from the Red Sea and Hong Kong that were long-term aquarium reared in separate facilities.

### Menthol bleaching seemingly led to stochastic changes in the microbiome of Galaxea

Menthol bleaching appeared to be associated with changes in the bacterial communities that differed between individual polyps and produced stochastic configurations. Unlike previous studies on Aiptasia that compared bleached and symbiotic individuals and found significantly different microbiomes (Röthig et al. 2016; Curtis et al. 2023), we could not identify a “signature of bleaching” as no bacterial taxa showed differential abundance between symbiotic states in our Galaxea colonies. This surprisingly included Symbiodiniaceae-associated bacteria that we were expecting to be reduced after the removal of Symbiodiniaceae (Fig. S5; Supplementary Materials and Methods). Such results could be an artifact of whole-tissue sampling in this study. As different coral compartments host distinct bacterial communities (Sweet et al. 2011; Ainsworth et al. 2015; Apprill et al. 2016), changes at the level of the gastroderm, where Symbiodiniaceae and their bacterial communities are located, might have been masked. Alternatively, it might indicate that Symbiodiniaceae-associated bacteria only repesent a small proportion of the community in Galaxea, or that they were able to persist in the absence of Symbiodiniaceae.

The lack of significant changes in bacterial taxa abundance between symbiotic and menthol-bleached polyps is surprising also given the known antimicrobial activity of menthol. Despite its effectiveness against various bacteria (i.e., *Escherichia coli, Staphylococcus aureus, Streptococcus* spp., *Lactobacillus* sp.) and its demonstrated selective activity on cigarette microbiomes (Trombetta et al. 2005; Chopyk et al. 2017; Mahzoon et al. 2022), direct impact of menthol on coral-associated bacteria may have been limited. Factors such as concentration, exposure time, and the varying susceptibility of bacterial species, alongside interactions with the coral environment and other microorganisms, likely contribute to the lack of an observable effect. These aspects, as well as further exploration of onset, duration, and specific mechanisms of action of menthol represent important areas for future research to elucidate its impact on coral-associated microbial communities.

A stochastic response to bleaching aligns well with the concept of an obligatory nature of the coral-algal symbiosis. In facultatively symbiotic cnidarians (such as the anemone Aiptasia), symbiotic and bleached can constitute alternative stable states, and it is thus possible to identify distinct and characteristic symbiotic and bleached microbiome configurations (Röthig et al. 2016; Curtis et al. 2023). Corals however depend on their algal symbiont for energy and other essential metabolic processes (Muscatine and Porter 1977; Muscatine 1990), and therefore the bleached state is not a stable alternative to the symbiotic state. In the bleached state the weakened coral host becomes progressively unable to regulate its microbial community, leaving room for the establishment of opportunistic bacteria, producing novel and stochastic combinations (Zaneveld et al. 2017).

The response of the bacterial microbiome to menthol bleaching seemed to differ between the two facilities. While we cannot draw definitive conclusions due to low replication, it is possible that differences in patterns of microbial response to menthol bleaching were linked to differences in rearing conditions. While these were largely replicated between facilities, feed type, tank volume and filtration systems differed. We therefore suggest that future studies incorporate an adequately replicated “facility” factor in their design, as well as food and seawater samples to better characterize these aspects.

### The microbiome of long-term aquarium-reared Galaxea fascicularis

The G. *fascicularis* polyps hosted simple microbiomes which were composed of a relatively small number of bacterial taxa (10-78 ASVs). While direct comparisons across studies that employed different PCR, sequencing, and analysis pipelines should generally be avoided, such numbers still appear small compared to previous characterizations of wild G. *fascicularis* from the South China Sea which reported 646-1,459 OTUs (Li et al. 2013) and compared to most other coral species, which typically harbor 100s to 1000s of bacterial taxa (e.g., Ziegler et al. 2016; Hernandez-Agreda et al. 2018; Pollock et al. 2018; Galand et al. 2023). We are unable to tell whether captivity caused a reduction in bacterial diversity as we lack direct comparison with the original wild colonies. However, we suspect that captivity favored a simplification of the microbiome, as stable and homogenous environmental conditions decrease both chances and need for the association with functionally and taxonomically diverse microbial partners. In fact, decreases in metabolic diversity and species richness have consistently been reported for tropical reef-building corals reared in closed systems (Kooperman et al. 2007; Vega Thurber et al. 2009; Pratte et al. 2015; Damjanovic et al. 2020), and for the anemone Aiptasia already after a few days of captivity (Hartman et al. 2020). Such effects could have been exacerbated by the use of filtered seawater during the bleaching phase, which largely reduced the pool of available microbes (Dungan et al. 2021b), and by the reduced structural complexity of the polyps that, compared to colonies, provide fewer micro-environments and ecological niches (Putnam et al. 2017; Morrow et al. 2022).

Regardless of the causes, simple (or simplified) microbiomes present the opportunity to identify essential associates and facilitate the development of microbial manipulation protocols to unravel holobiont functioning (Jaspers et al. 2019; Puntin et al. 2022b). Culturing corals in sterile seawater may help to limit the horizontal acquisition of transient microbes and thus favor proliferation of core or stable members for detailed characterization (Dungan et al. 2021b). Also, a simple microbiome facilitates the elimination of bacterial populations to produce gnotobiotic or axenic hosts, which could subsequently be re-inoculated to produce a range of host-bacteria combinations to test microbial functions and inter-partner dynamics (Fraune et al. 2015; Murillo-Rincon et al. 2017; Jaspers et al. 2019; Taubenheim et al. 2020). Reduced microbial complexity—whether due to captivity or other factors—might therefore provide advantages for these specific experimental approaches with the Galaxea model.

Interestingly, the coral colonies tested here maintained distinct bacterial microbiomes even after long-term co-culturing, which supports a degree of host genotype effects controlling the microbiome composition, as previously reported from Hydrozoan and other coral species in the field (Pollock et al. 2018; Dubé et al. 2021). Surprisingly, the microbiome of one Red Sea colony was highly similar to that of Hong Kong colonies. This appears counterintuitive as colonies from the Red Sea and Hong Kong may belong to different Galaxea lineages (sensu Wepfer et al. 2020), and considering the large differences in environmental conditions at their origin. While these colonies were also maintained in separate facilities, rearing conditions were similar at both locations (i.e., temperature, salinity, illumination) and may have induced convergence of microbial community composition (Dubé et al. 2021). On the other hand, G. *fascicularis* is a polyphyletic species that contains several morphologically cryptic lineages (Wepfer et al. 2020) and the dissimilarity between colonies from the Red Sea could reflect host phylogentic differences (i.e., cryptic species within the same region).

Besides host genotype, Symbiodiniaceae community composition could also explain differences in bacterial community composition between colonies (Littman et al. 2010; Bernasconi et al. 2019). To investigate this aspect, we also characterized the Symbiodiniacea communities of the same polyps (Supplementary Materials and Methods). While only a small proportion of samples were successfully sequenced, we noticed emerging patterns of Symbiodiniaceae-bacteria co-occurrence that warrant further investigation (Fig. S5). We therefore recommend that future studies characterize a larger number of Galaxea holobionts at multiple locations across the species distribution range to explore links between host-Symbiodiniaceae-bacteria associations. This could elucidate the influence of each member on coral holobiont compositions and functioning.

### Core bacterial associates of Galaxea

We defined core or stable microbial associates based on prevalence of ASVs across treatment groups and polyps. Notably, no taxa was present in 100 % of the samples, suggesting a certain degree of microbiome variability within this coral host species. Among the 28 core ASVs (occuring in all groups), the five most frequent and abundant were assigned to the genera *Alteromonas* and *Ruegeria*. Both *Alteromonas* and *Ruegeria* are common coral-associates reported from at least 20 other coral species, and sequences assigned to these two genera ranked 6^th^ and 33^rd^ most abundant in the Coral Microbial Database (Huggett and Apprill 2019).

*Ruegeria* spp. are commonly and consistently found in association with G. *fascicularis* in wild and aquarium-reared colonies, from Hong Kong to Japan and across seasons (Cai et al. 2018a, 2018b; Miura et al. 2019; Tang et al. 2020; Kitamura et al. 2021). Indeed, the two *Ruegeria* core ASVs had identical sequences with *Ruegeria* from G. *fascicularis* from Hainan and Japan that were maintained under aquarium conditions comparable to ours (Miura et al. 2019; Zhou et al. 2020). This shows that the Galaxea-*Ruegeria* association is highly conserved and therefore putatively biologically relevant. *Ruegeria* strains isolated from G. *fascicularis* were indeed previously identified as potential probiotics, through inhibitory activity towards the coral pathogen Vibrio *coralliilyticus* (Kitamura et al. 2021). The ubiquitous and persistent Galaxea-*Ruegeria* association thus warrants attention in future investigations.

The potential role of *Alteromonas* spp. in the Galaxea holobiont functioning also deserves attention. Despite their high abundance and prevalence in this study (and in corals in general), the role of *Alteromonas* spp. remains controversial. They have been considered pathogenic, owing to their co-occurrence with coral diseases (e.g., Sunagawa et al. 2009), but also listed as candidate probiotic for their free radical scavenging abilities (Raina et al. 2009; Dungan et al. 2021a), which could be linked to their consistent association with Symbiodiniaceae (Lawson et al. 2018; Nitschke et al. 2020).

Bacteria in the family *Endozoicomonadaceae* are the most prominent members of coral microbiomes in a range of coral species (Morrow et al. 2012; Bayer et al. 2013; Neave et al. 2017; Pogoreutz et al. 2022), which have been investigated for their involvement in holobiont metabolism, for example in the C and S cycles (Neave et al. 2016; Ide et al. 2022). Yet, *Endozoicomonadaceae* were only present in approximately half of Galaxea polyps, at between 0.06 and 39.12 % (mean 5.78 %) relative abundance. This is slightly lower than in wild G. *fascicularis* from Hong Kong (< 10 % relative abundance) in which, however, *Endozoicomonas* spp. were present in all samples (Cai et al. 2018a, 2018b). Such scarcity and inconsistent presence therefore suggests that *Endozoicomonas* spp. might not play an essential role in the G. *fascicularis* under captive conditions. If further proven, a lack of reliance on *Endozoicomonas* spp. in captivity could offer insights into which functions benefit from microbial help in the wild, hence highlight the role of this bacterial associate in the coral holobiont in the wider context.

## Conclusions

Model organisms provide powerful tools for unraveling holobiont complexity. These models can be used to test hypotheses of functional relationships and inter-partner interactions through holobiont manipulation. To complement current cnidarian model systems such as Aiptasia and Hydra, we recently proposed the adoption of Galaxea fascicularis as a true coral model owing to its suitability to aquarium rearing and reproduction, and manipulation of its association with Symbiodiniaceae following menthol bleaching (Puntin et al. 2022a). However, how this bleaching treatment affected the bacterial microbiome remained to be explored.

In this study, we provided the first baseline assessment of the response of the Galaxea bacterial microbiome to menthol bleaching. Overall, menthol bleaching appeared to induce stochastic changes in the microbiome seemingly indicating dysbiosis. The response of the bacterial microbiome to bleaching differed in trend between the two facilities, possibly reflecting differences in rearing conditions, which remain to be addressed. Despite co-culturing in shared aquaria, symbiotic polyps originating from different colonies maintained distinct community assemblies. This suggest links to host and/or Symbiodiniaceae identity that warrant further investigation.

Bacterial communities of the captive Galaxea colonies were composed of a small number of taxa. A simple microbiome could facilitate both characterization and experimental manipulation, and guide the identification of essential (“core”) members among the retained associates. In this regard, we identified Ruegeria spp. and *Alteromonas* spp. as candidate associates for further functional interrogation. In conclusion, our study contributes valuable information towards a better characterization of the Galaxea holobiont, as well as its continued development and establishment as a coral model system.

## Acknowledgements

We thank André Dietzmann, Kara E. Engelhardt, Patrick Schubert, and Christina Anding (all JLU) for assistance during experiments; and Talia Morris and Hugh Spencer (Austrop) for offering their office and kindness, allowing GP to work on this manuscript from a very special place.

## Data, scripts, code, and supplementary information availability

Data are available online:

Raw sequencing data submitted to NCBI SRA BioProject PRJNA947274

Representative sequences (ASVs) submitted to NCBI GenBank Accession numbers OQ677536 to OQ677992

Additional data, scripts, and code are available online: https://doi.org/10.5281/zenodo.10551928 (Puntin 2024)

