## Supplementary Figures, Tables, and Materials and Methods for "The bacterial microbiome of symbiotic and menthol-bleached polyps of long-term aquarium-reared *Galaxea fascicularis*" (from another version of this preprint: version 4, 2024-05-21)

Figure S1

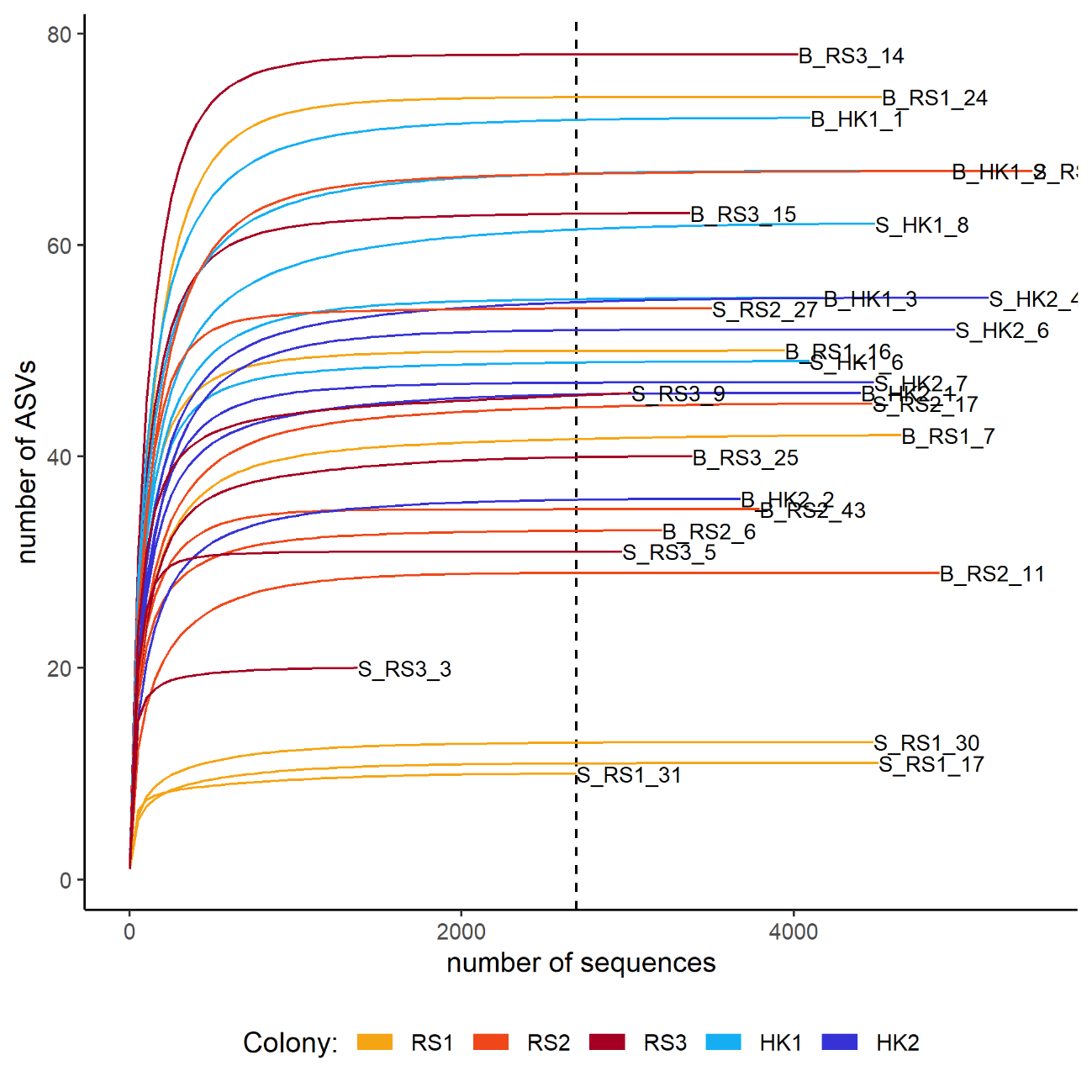

**Figure** **S1** - Rarefaction curve. Dashed line indicates rarefaction depth applied (2,690).

Figure S2

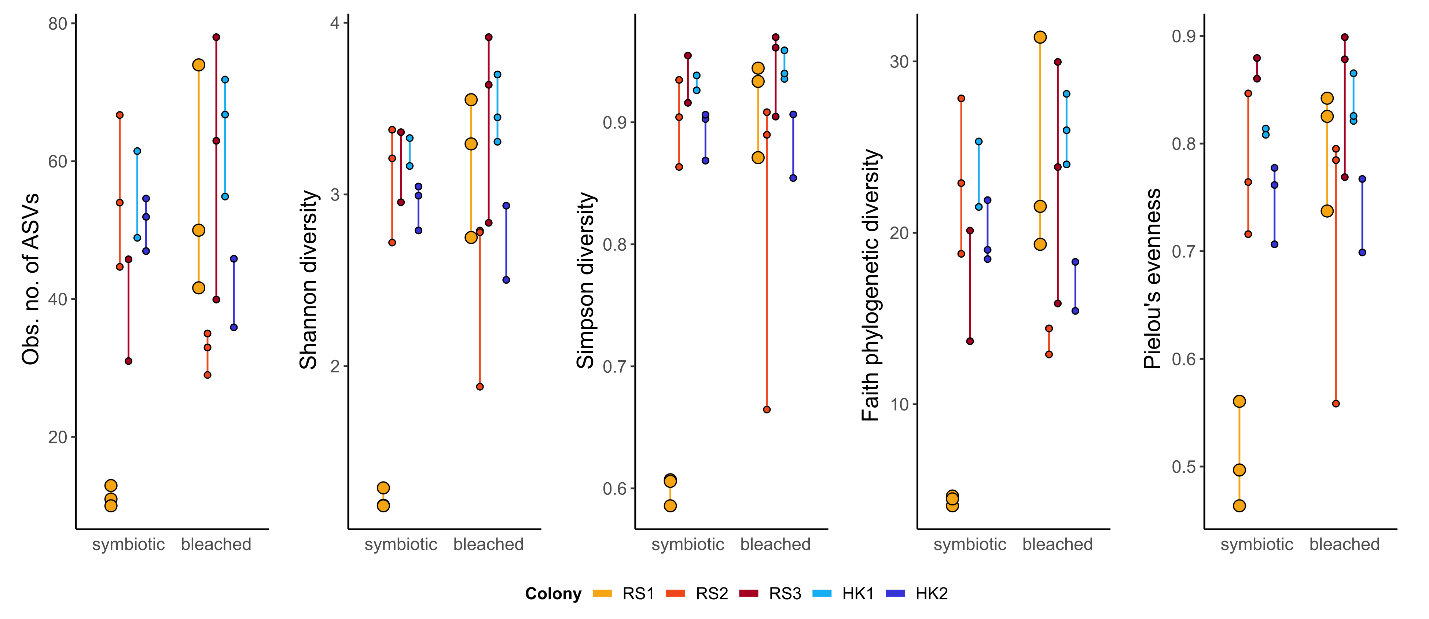

**Figure** **S2** - Comparison of microbial diversity and richness between symbiotic and bleached colonies, displayed as raw data points and ranges. Differences between symbiotic and bleached polyps could only be tested for RS1 and RS2 due to inadequate replication in the remaining colonies (results in Tab. S3). The difference between symbiotic states is significant for all tested indices in colony RS1, which is highlighted here with larger points size for ease of identification.

Figure S3

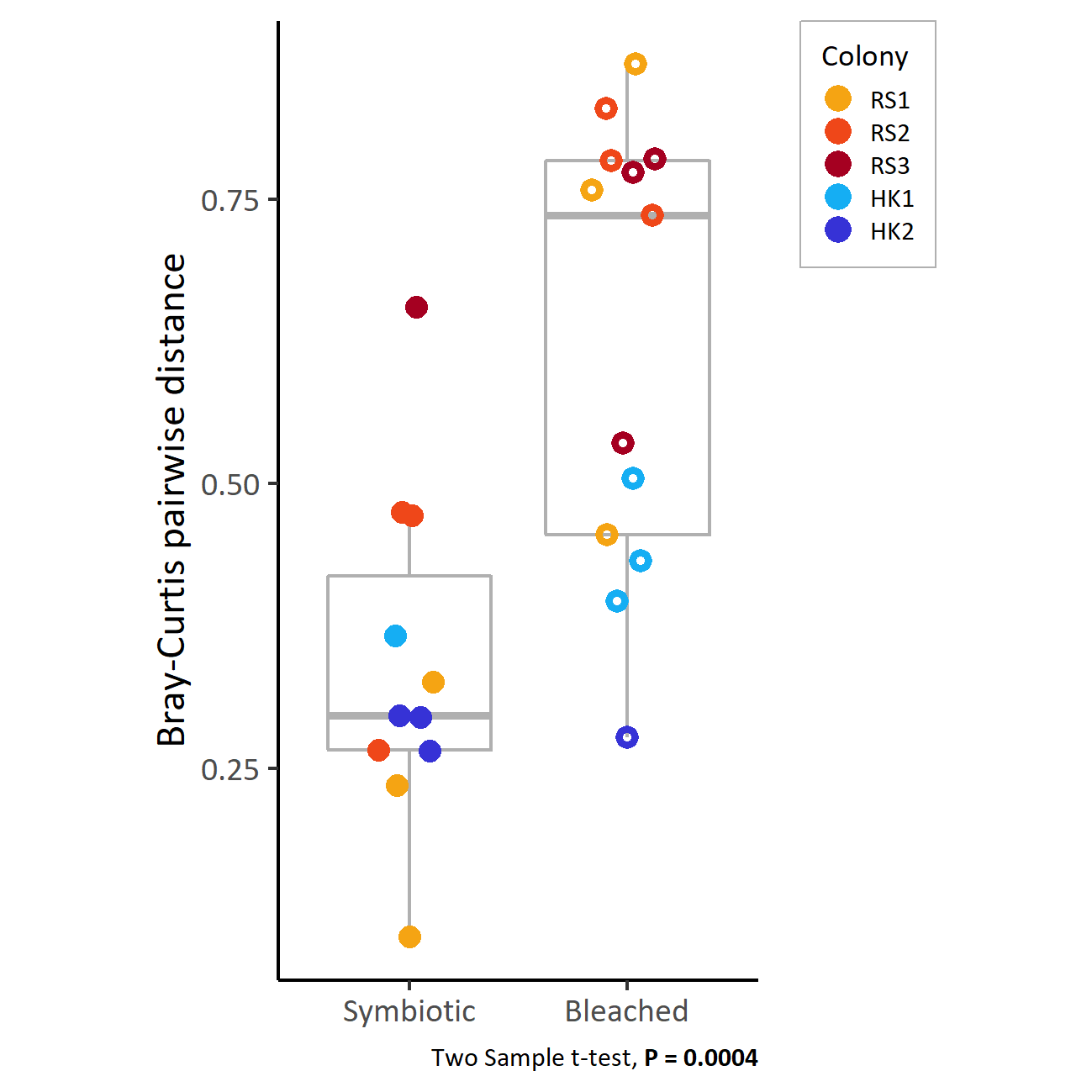

**Figure S3** - Comparison of (within group) dissimilarity between symbiotic and bleached samples. Dissimilarity as pairwise distance (Bray-Curtis) between samples of the same colony, by symbiotic state. All groups considered (i.e., with *n* = 2 and *n* = 3).

Figure S4

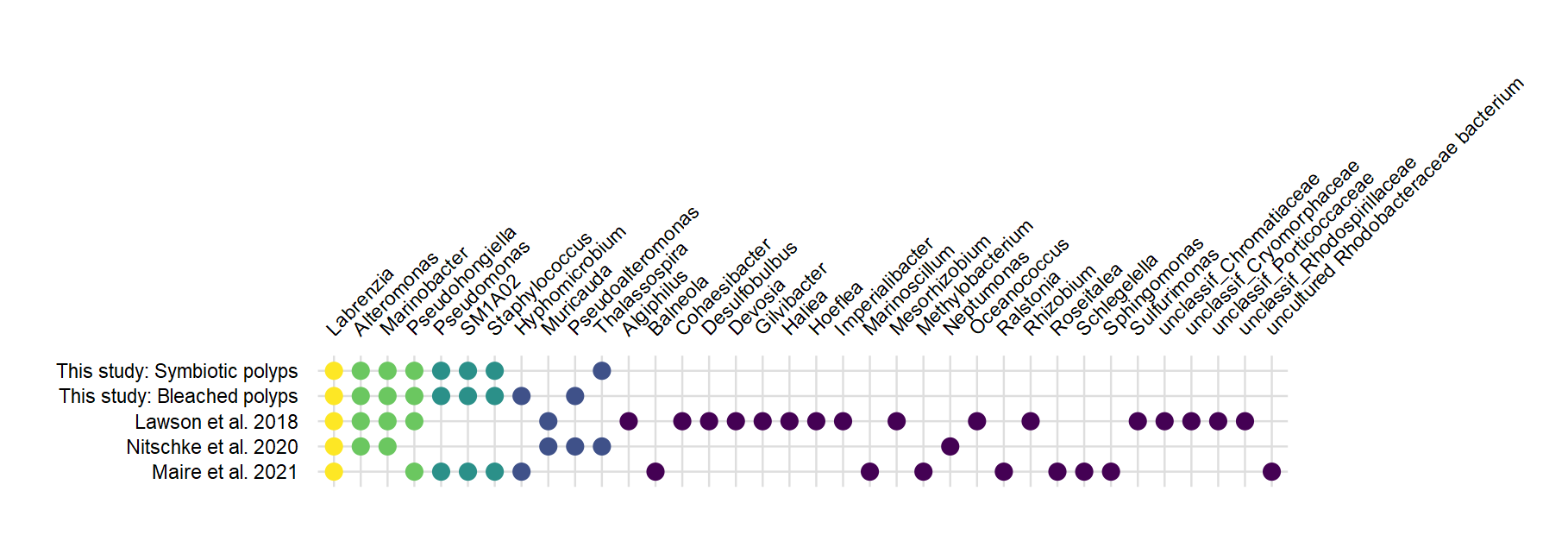

**Figure S4** - Comparison between our findings (“This study”) and other studies on Symbiodiniaceae-associated bacteria displayed as presence/absence of bacteria genera. All genera reported as associated with Symbiodiniaceae in any of the reference studies (Lawson et al. 2018, Nitschke et al. 2020, and Maire et al. 2021) are displayed, but genera from this study that had no match with any of these three studies are omitted from this plot. Points are color-coded by number of intersections (co-occurrences). For details regarding the studies considered, with specification of the respective data, please refer to the Supplementary Materials and Methods section.

Figure S5

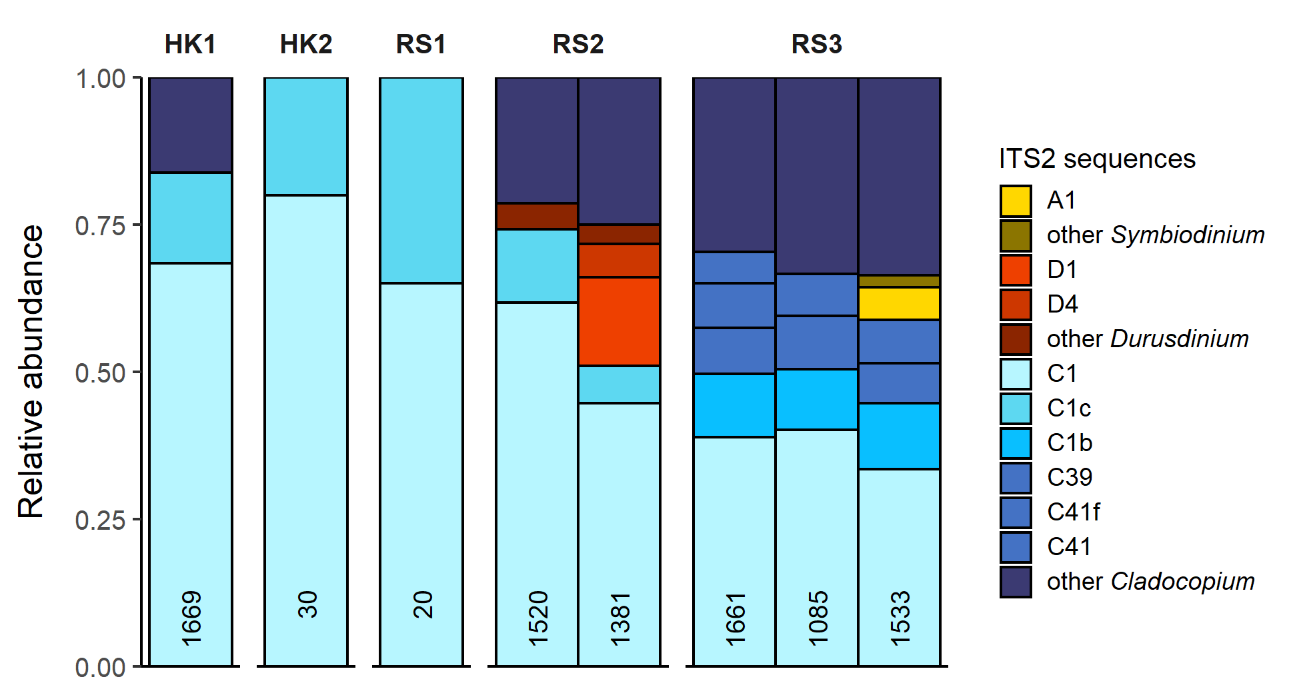

**Figure S5** - Relative abundance of Symbiodiniaceae ITS2 sequences by individual symbiotic sample. Based on SymPortal (post-MED) ITS2 output. Sequences with relative abundances < 5 % (by sample) are grouped per genus as “other *”. The total number of sequences per sample is indicated at the base of each column.

Supplementary Tables

Table S1

**Table S1** - Statistical testing of difference in community diversity and evenness between symbiotic (n = 13) and bleached (n = 14) colonies. The choice of the statistical test followed data inspection, where Shannon diversity, Simpson evenness, and Pielou’s evenness showed non-normal distribution (qq-plots), while Simpson evenness also showed unequal variance (F-test, Bartlett test).

| **Alpha diversity metric** | **Test** | **Statistic** | **P** |
| --- | --- | --- | --- |
| Observed richness | unpaired 2-samples t-test | 1.417457 | 0.0843 |
| Shannon diversity | unpaired 2-samples t-test | 1.582024 | 0.0631 |
| Simpson evenness | Welch t-test | 1.45282 | 0.0816 |
| Faith phylogenetic diversity | unpaired 2-samples t-test | 1.446646 | 0.0802 |
| Pielou's evenness | unpaired 2-samples t-test | 1.44496 | 0.0804 |
| Chao1 | unpaired 2-samples t-test | 1.396447 | 0.0874 |
| Shannon diversity | Mann-Whitney U-test | 113 | 0.1510 |
| Simpson evenness | Mann-Whitney U-test | 123 | 0.0639 |
| Pielou's evenness | Mann-Whitney U-test | 117 | 0.1100 |

Table S2

**Table 2** - Summary of alpha diversity values by colony and symbiotic state.

| **State** | **Origin** | **Colony** | **n** | **Observed** | **Chao1** | **Shannon** | **Simpson** | **Pielou** | **Faith_PD** |
| --- | --- | --- | --- | --- | --- | --- | --- | --- | --- |
| Symbiotic | Red Sea | RS1 | 3 | 11.301 | 11.302 | 1.222 | 0.600 | 0.507 | 4.394 |
| Symbiotic | Red Sea | RS2 | 3 | 55.119 | 55.268 | 3.102 | 0.901 | 0.776 | 23.172 |
| Symbiotic | Red Sea | RS3 | 2 | 38.386 | 38.758 | 3.159 | 0.935 | 0.870 | 16.902 |
| Symbiotic | Hong Kong | HK1 | 2 | 55.174 | 55.519 | 3.247 | 0.932 | 0.811 | 23.419 |
| Symbiotic | Hong Kong | HK2 | 3 | 51.158 | 51.310 | 2.943 | 0.892 | 0.748 | 19.793 |
| Bleached | Red Sea | RS1 | 3 | 55.195 | 55.279 | 3.199 | 0.916 | 0.802 | 24.095 |
| Bleached | Red Sea | RS2 | 3 | 32.320 | 32.322 | 2.483 | 0.821 | 0.713 | 13.914 |
| Bleached | Red Sea | RS3 | 3 | 60.292 | 60.327 | 3.463 | 0.945 | 0.849 | 23.227 |
| Bleached | Hong Kong | HK1 | 3 | 64.491 | 64.623 | 3.485 | 0.945 | 0.837 | 26.028 |
| Bleached | Hong Kong | HK2 | 2 | 40.869 | 40.967 | 2.718 | 0.880 | 0.733 | 16.872 |

Table S3

**Table S3** - Statistical testing of difference in community diversity and evenness between symbiotic (group2, n2) and bleached (group1, n1) polyps using the unequal variances unpaired t-test (Welch’s test). Testing was limited to colonies RS1 and RS2 due to insufficient replication in the remaining colonies (n1 or n2 < 3 ).

| **Colony** | **Alpha diversity index** | **Group1** | **Group2** | **n1** | **n2** | **Statistic** | **df** | **P** |  |
| --- | --- | --- | --- | --- | --- | --- | --- | --- | --- |
| RS1 | Observed | bleached | symbiotic | 3 | 3 | 4.51 | 2.03 | 0.0445 | * |
| RS1 | Shannon | bleached | symbiotic | 3 | 3 | 8.27 | 2.08 | 0.0127 | * |
| RS1 | Simpson | bleached | symbiotic | 3 | 3 | 13.27 | 2.36 | 0.0028 | ** |
| RS1 | Pielou | bleached | symbiotic | 3 | 3 | 6.82 | 3.93 | 0.0026 | ** |
| RS1 | Faith_PD | bleached | symbiotic | 3 | 3 | 5.30 | 2.01 | 0.0335 | * |
| RS2 | Observed | bleached | symbiotic | 3 | 3 | -3.44 | 2.30 | 0.0613 |  |
| RS2 | Shannon | bleached | symbiotic | 3 | 3 | -1.72 | 3.45 | 0.1720 |  |
| RS2 | Simpson | bleached | symbiotic | 3 | 3 | -0.99 | 2.28 | 0.4170 |  |
| RS2 | Pielou | bleached | symbiotic | 3 | 3 | -0.73 | 2.92 | 0.5200 |  |
| RS2 | Faith_PD | bleached | symbiotic | 3 | 3 | -3.47 | 2.15 | 0.0668 |  |

Table S4

**Table S4** - Summary of taxonomy, abundance, and prevalence across the data set for the 28 core ASVs that occur across all groups (*state* × *origin*). ‘Overall’ (n = 28), ‘Symbiotic’ (n = 14), ‘Bleached’ (n = 14). Taxonomy is reported at the genus level unless otherwise indicated in parenthesis ("C" = Class, “F” = Family). Blocks indicate the 70 % and 60 % cut-offs of overall abundance across samples. ASV ids are ordered by their abundance across the whole data set. Note that the eight most abundant ASVs reported in this table are also the eight most abundant overall (regardless of which *state* × *origin* groups are considered). For the full table (515 ASVs, non-rarefied data) see: <https://doi.org/10.5281/zenodo.10551928> (~/out/Gfas_16S/core_mb/nonrarefied/ASV_occurrence_summary_all.csv)

|  |  | **No. of samples found in** | | | **% of samples found in** | | | **abundance** | | | **abundance** | | |
| --- | --- | --- | --- | --- | --- | --- | --- | --- | --- | --- | --- | --- | --- |
|  |  |  |  |  |  |  |  | **(reads count)** | | | **(% of all reads)** | | |
| **Taxonomy** | **ASV id** | **Overall** | **Symbiotic** | **Bleached** | **Overall** | **Symbiotic** | **Bleached** | **Overall** | **Symbiotic** | **Bleached** | **Overall** | **Symbiotic** | **Bleached** |
| **(lowest taxonomic level when above genus)** |  |  |  |  |  |  |  |  |  |  |  |  |  |
| *Alteromonas* | ASV_003 | 25 | 12 | 13 | 89.3 | 85.7 | 92.9 | 9281 | 5989 | 3292 | 8.2 | 10.8 | 5.8 |
| *Ruegeria* | ASV_001 | 22 | 11 | 11 | 78.6 | 78.6 | 78.6 | 13836 | 8117 | 5719 | 12.3 | 14.6 | 10.0 |
| *Alteromonas* | ASV_008 | 20 | 8 | 12 | 71.4 | 57.1 | 85.7 | 2838 | 1365 | 1473 | 2.5 | 2.5 | 2.6 |
| *Nautella* | ASV_005 | 19 | 9 | 10 | 67.9 | 64.3 | 71.4 | 3883 | 2888 | 995 | 3.4 | 5.2 | 1.7 |
| *Ruegeria* | ASV_006 | 19 | 9 | 10 | 67.9 | 64.3 | 71.4 | 3523 | 1668 | 1855 | 3.1 | 3.0 | 3.2 |
| *Gammaproteobacteria* (C) | ASV_017 | 18 | 8 | 10 | 64.3 | 57.1 | 71.4 | 1130 | 503 | 627 | 1.0 | 0.9 | 1.1 |
| *Cryomorphaceae* (F) | ASV_018 | 17 | 8 | 9 | 60.7 | 57.1 | 64.3 | 1114 | 659 | 455 | 1.0 | 1.2 | 0.8 |
| *Hyphomonadaceae* (F) | ASV_020 | 17 | 6 | 11 | 60.7 | 42.9 | 78.6 | 951 | 174 | 777 | 0.8 | 0.3 | 1.4 |
| *Micavibrionaceae* (F) | ASV_016 | 16 | 9 | 7 | 57.1 | 64.3 | 50.0 | 1150 | 638 | 512 | 1.0 | 1.1 | 0.9 |
| *Methylotenera* | ASV_028 | 16 | 8 | 8 | 57.1 | 57.1 | 57.1 | 644 | 286 | 358 | 0.6 | 0.5 | 0.6 |
| *Oleibacter* | ASV_013 | 15 | 7 | 8 | 53.6 | 50.0 | 57.1 | 1229 | 465 | 764 | 1.1 | 0.8 | 1.3 |
| *Colwelliaceae* (F) | ASV_014 | 15 | 6 | 9 | 53.6 | 42.9 | 64.3 | 1225 | 532 | 693 | 1.1 | 1.0 | 1.2 |
| *Acinetobacter* | ASV_011 | 13 | 7 | 6 | 46.4 | 50.0 | 42.9 | 1679 | 1214 | 465 | 1.5 | 2.2 | 0.8 |
| *Rhodobacteraceae* (F) | ASV_025 | 13 | 7 | 6 | 46.4 | 50.0 | 42.9 | 759 | 560 | 199 | 0.7 | 1.0 | 0.3 |
| *Peredibacter* | ASV_041 | 13 | 7 | 6 | 46.4 | 50.0 | 42.9 | 494 | 396 | 98 | 0.4 | 0.7 | 0.2 |
| *HTCC5015*  (F: Arenicellaceae) | ASV_042 | 13 | 4 | 9 | 46.4 | 28.6 | 64.3 | 490 | 89 | 401 | 0.4 | 0.2 | 0.7 |
| *Algimonas* | ASV_031 | 12 | 4 | 8 | 42.9 | 28.6 | 57.1 | 613 | 195 | 418 | 0.5 | 0.4 | 0.7 |
| *HIMB11*  (F:Rhodobacteraceae) | ASV_022 | 11 | 3 | 8 | 39.3 | 21.4 | 57.1 | 902 | 152 | 750 | 0.8 | 0.3 | 1.3 |
| *Ruegeria* | ASV_021 | 10 | 7 | 3 | 35.7 | 50.0 | 21.4 | 945 | 762 | 183 | 0.8 | 1.4 | 0.3 |
| *Vibrio* | ASV_034 | 10 | 4 | 6 | 35.7 | 28.6 | 42.9 | 584 | 260 | 324 | 0.5 | 0.5 | 0.6 |
| *Flavobacteriaceae* (F) | ASV_038 | 10 | 4 | 6 | 35.7 | 28.6 | 42.9 | 507 | 204 | 303 | 0.4 | 0.4 | 0.5 |
| *Labrenzia* | ASV_055 | 10 | 3 | 7 | 35.7 | 21.4 | 50.0 | 359 | 68 | 291 | 0.3 | 0.1 | 0.5 |
| *Alteromonas* | ASV_033 | 9 | 5 | 4 | 32.1 | 35.7 | 28.6 | 587 | 395 | 192 | 0.5 | 0.7 | 0.3 |
| *Micavibrionaceae* (F) | ASV_054 | 8 | 4 | 4 | 28.6 | 28.6 | 28.6 | 382 | 202 | 180 | 0.3 | 0.4 | 0.3 |
| *Sphingomonadaceae* (F) | ASV_050 | 7 | 3 | 4 | 25.0 | 21.4 | 28.6 | 404 | 235 | 169 | 0.4 | 0.4 | 0.3 |
| *Gammaproteobacteria* (C) | ASV_059 | 7 | 4 | 3 | 25.0 | 28.6 | 21.4 | 293 | 177 | 116 | 0.3 | 0.3 | 0.2 |
| *Lewinella* | ASV_062 | 6 | 2 | 4 | 21.4 | 14.3 | 28.6 | 260 | 46 | 214 | 0.2 | 0.1 | 0.4 |
| *Arenicella* | ASV_117 | 5 | 3 | 2 | 17.9 | 21.4 | 14.3 | 89 | 69 | 20 | 0.1 | 0.1 | 0.0 |

Supplementary Materials and Methods

Symbiodiniaceae-associated bacterial taxa from other studies

We identified studies that characterized the bacterial communities associated with Symbiodiniaceae and compared these with the bacterial taxa found in our study, in either symbiotic or aposymbiotic (menthol bleached) samples. This way, we could preliminarily inspect for patterns of presence/absence of taxa between symbiotic states, on the assumption that Symbiodiniaceae-associated bacteria should be present in symbiotic corals and absent in bleached corals. This approach produced Figure S4 and incorporated the studies by Lawson et al. (2018), Nitschke et al. (2020), and Maire et al. (2021).

For more details, the formatted data and R scripts are available at: <https://doi.org/10.5281/zenodo.10551928>

Lawson et al. 2018

Study summary

Characterized the bacterial communities associated with Symbiodiniaceae cultures, spanning 18 cultures across 5 genera (former clades) to define the core Symbiodiniaceae microbiome. Identified three OTUs, which were present in all cultures and corresponded to *Labrenzia*, *Marinobacter,* and *Chromatiaceae*.

Data considered

Supplementary material, Table S3, with description: **“**Operational taxonomic units (OTUs) defined as core members of the bacterial communities of *Symbiodinium* cultures and their corresponding GenBank accession numbers.” This table reports the core taxa/ASVs overall (across all Symbiodiniaceae genera) and for each genus separately.

Extracted information and use

Information on bacteria taxonomy was extracted (from the article’s Table S3) from the column “Taxonomic ID”, which reports the bacterial taxonomy to the genus level when available, else to higher level preceded by “UC” (for “unclassified”). "UC " was replaced with "unclassif_" to match the naming system adopted in our study, and screened for matches. This resulted in four matches, all at the genus level.

Nitschke et al. 2020

Study summary

Study on symbiolites (photosynthesis-induced microbialites formed by calcifying co-cultures of Symbiodiniaceae and bacteria), with comparison of bacterial communities of symbiolites-producing (SP) and non-symbiolites producing (NP) Symbiodiniaceae cultures.

Data considered

Manuscript Table 2: “Bacterial isolates from symbiodiniacean cultures”, with caption: “List of bacterial strains used in this study, including their taxonomic affiliation, GenBank accession numbers, and the Symbiodiniaceae strain of origin (ITS2 type). […]”.

Bacteria were isolated from Symbiodiniaceae cultures belonging to strains (ITS2 types): A1, A2 (*Symbiodinium*), B1 (*Breviolum*), and C2 (*Cladocopium*).

Extracted information and use

The table reports taxonomy down to the species level (for all isolates). Genus level affiliation was extracted and, together with species names, cross checked with the data from our study. This resulted in 5 matches at the genus level and no matches at the species level.

Maire et al. 2021

Study summary

Characterized the bacterial communities associated with 11 Symbiodiniaceae strains spanning nine species and six genera, and distinguishing between intracellular, closely associated (on Symbiodiniaceae outer cell surface), and loosely associated bacterial communities.

Data considered

Supplementary “Dataset S3” corresponding to file “41396_2021_902_MOESM4_ESM.xlsx”, with description: “Relative abundances of intracellular (A), closely associated (B), and loosely associated (C) core genera in all Symbiodiniaceae samples. A core genus is a genus that is present in every Symbiodiniaceae species within a given location.”, which contains the phylogeny and abundance of the core genera for each location (“intracellular”, “closely associated” and “loosely associated”) for all 11 strains. With “core genera” corresponding to the genera found in all 11 Symbiodiniaceae strains for each location. The 11 Symbiodiniaceae strains (listed in Table S1, file “41396_2021_902_MOESM3_ESM.xlsx”) belonged to the genera Symbiodinium, Breviolum, Cladocopium, Durusdinium, Fugacium, Gerakladium.

Extracted information and use

The names of the “core genera” for all locations (all three sheets: “A - Intracellular core genera”, “B - Closely-assoc core genera”, and “C - Loosely-assoc core genera”) were extracted and crossed checked with the data from our study, which resulted in 6 matches.

Characterization of Symbiodiniaceae communities of *G. fascicularis* polyps

Sampling and Symbiodiniaceae community analysis

The characterization of Symbiodiniaceae communities was performed using the same polyps’ DNA extract employed for the 16S rRNA amplicon sequencing (characterization of bacterial communities) and sequenced in the same sequencing run (see main manuscript for details).

Symbiodiniaceae ITS2 region was amplified using the primers SYM_VAR_5.8S2 and SYM_VAR (Hume et al. 2018), and raw ITS2 sequencing data were analyzed using the SymPortal workflow remote instance (Hume et al. 2019). ITS2 sequencing produced poor results and it was only possible to characterize eight of the symbiotic samples (6 Red Sea, 2 Hong Kong; see Fig. 1). Of these, two did not pass quality check (i.e., they had < 200 sequences/genus) and SymPortal could not predict ITS2 type profiles. Therefore, we report (post-MED) ITS2 sequences.

Results

All colonies of *Galaxea fascicularis* were dominated by *Cladocopium* spp. symbionts. Symbiodiniaceae composition was consistent in polyps from each colony and differed by region. *Cladocopium* spp. ITS2 sequences accounted for > 92 % of the sequences in all samples but one (RS2, 76 %) (Fig. S5). Of these, C1 was by large the dominant ITS2 sequence (~ 33 – 80 % relative abundance). Sequences C1b, C41, and C41f were exclusively and consistently found in polyps of one Red Sea colony (RS3), where they collectively accounted for ~ 26 % of reads. *Durusdinium* spp. sequences were found in only one colony from the Red Sea (RS2), where they accounted for 4 % and 24 % of the sequences, and of which the most abundant sequences were D1 and D4 (Fig. S5). One Red Sea sample (RS3) also hosted sequence A1 (genus *Symbiodinium*) at ~ 5 % abundance. In both Hong Kong colonies C1 was the dominant sequence (68 – 80 %), with C1c present in lower abundance (15 – 20 %) (Fig. S5).
